# Quantification of transcript isoforms at the single-cell level using SCALPEL

**DOI:** 10.1101/2024.06.21.600022

**Authors:** Franz Ake, Sandra M. Fernández-Moya, Marcel Schilling, Akshay Jaya Ganesh, Ana Gutiérrez-Franco, Lei Li, Mireya Plass

**Affiliations:** Gene Regulation of Cell Identity, Regenerative Medicine Program, Bellvitge Institute for Biomedical Research (IDIBELL), 08908, L’Hospitalet del Llobregat, Spain; Program for Advancing Clinical Translation of Regenerative Medicine of Catalonia, P-CMR[C], 08908, L’Hospitalet del Llobregat, Spain; Institute of Systems and Physical Biology, Shenzhen Bay Laboratory, Shenzhen, 518055, China; Center for Networked Biomedical Research on Bioengineering, Biomaterials and Nanomedicine (CIBER-BBN), 28029, Madrid, Spain

## Abstract

Single-cell RNA sequencing (scRNA-seq) has facilitated the study of gene expression and the development of new tools to quantify transcript in individual cells. Yet, most of these methods have low sensitivity and accuracy. Here we present SCALPEL, a Nextflow-based tool to quantify and characterize transcript isoforms at the single-cell level using standard 3’ based scRNA-seq data. SCALPEL predictions have higher sensitivity than other tools and can be validated experimentally. We have used SCALPEL to study the changes in isoform usage during mouse spermatogenesis and in the differentiation of induced pluripotent stem cells (iPSCs) to neural progenitors. These analyses allow the identification of novel cell populations that cannot be defined using conventional gene expression profiles, confirm known changes in 3’ UTR length during cell differentiation, and identify cell-type specific miRNA signatures controlling isoform expression in individual cells. Together, our work highlights how SCALPEL expands the current scRNA-seq toolset to explore post-transcriptional gene regulation in individual cells from different species, tissues, and technologies to investigate the variability and the specificity of gene regulatory mechanisms at the single-cell level.

## Main

Alternative polyadenylation (APA) is a general mechanism of post-transcriptional regulation that significantly contributes to the diversification of gene expression patterns under diverse physiological and pathological conditions^1^. APA defines the end of transcripts by selecting one of the available polyA sites (PAS) at the 3’ end of genes, resulting in the generation of multiple mature RNA isoforms from the same pre-mRNA^2^. These isoforms may have different coding regions or contain distinct 3’ untranslated regions (3’ UTRs), which contain regulatory elements influencing mRNA stability, localization, and translational efficiency^3–5^. Transcriptomic studies have demonstrated that APA is highly regulated in a tissue specific manner^6^ and plays a crucial role in various biological processes, including cellular differentiation^7^, development^8–10^, and response to environmental cues^11^. Alterations in APA patterns have been linked to various diseases, where they can lead to aberrant gene expression and even cancer^12,13^.

The development of high-throughput single-cell transcriptomics technologies (scRNA-seq) has led to the emergence of computational methods to characterize the transcriptomic profile of thousands of individual cells in a single experiment^14^. While these methods are mainly used to quantify gene expression, 3’ tag-based scRNA-seq protocols such as Drop-seq^15^ or 10x genomics provide opportunities to study 3’ end isoform diversity. Currently, only a few computational tools allow to study isoform diversity generated by APA in scRNA-seq data and most of them face significant drawbacks. They often fail to detect polyadenylation sites (PAS) with low read coverage due to the sparse nature of single-cell data and they lack the precision needed to accurately pinpoint the exact PAS locations, leading to potential misidentification and incomplete characterization of isoform diversity^16–19^. Alternative methods based on isoform quantification such as scUTRquant^20^ have been shown to be more powerful in quantifying transcript diversity from scRNA-seq data. Yet, the main power of this method relies on an improved curated 3’ end annotation that is not available for most species.

Here, we present SCALPEL, a Nextflow workflow^21^ to quantify isoform expression using commonly used 3’ tag-based scRNA-seq data. SCALPEL workflow is divided into three main modules (Fig 1a). In the first module, raw sequencing data and annotation files are processed to perform bulk quantification of the annotated isoforms. These isoforms are then truncated and collapsed, giving rise to a set of distinct isoforms with different 3’ ends for quantification at single-cell resolution. In the second module, scRNA-seq reads are mapped on the set of selected isoforms and reads coming from pre-mRNAs or resulting from internal priming (IP) events are discarded. In the last module, isoforms are quantified in individual cells and an isoform digital gene expression matrix (iDGE) is generated (Fig. S1). The iDGE can be processed to perform downstream single-cell level analyses such as dimensionality reduction, clustering, marker discovery and trajectory inference. Furthermore, it can also be used to study differential isoform usage (DIU) and visualize isoform coverage using additional functions included in SCALPEL repository.

**Figure 1:**
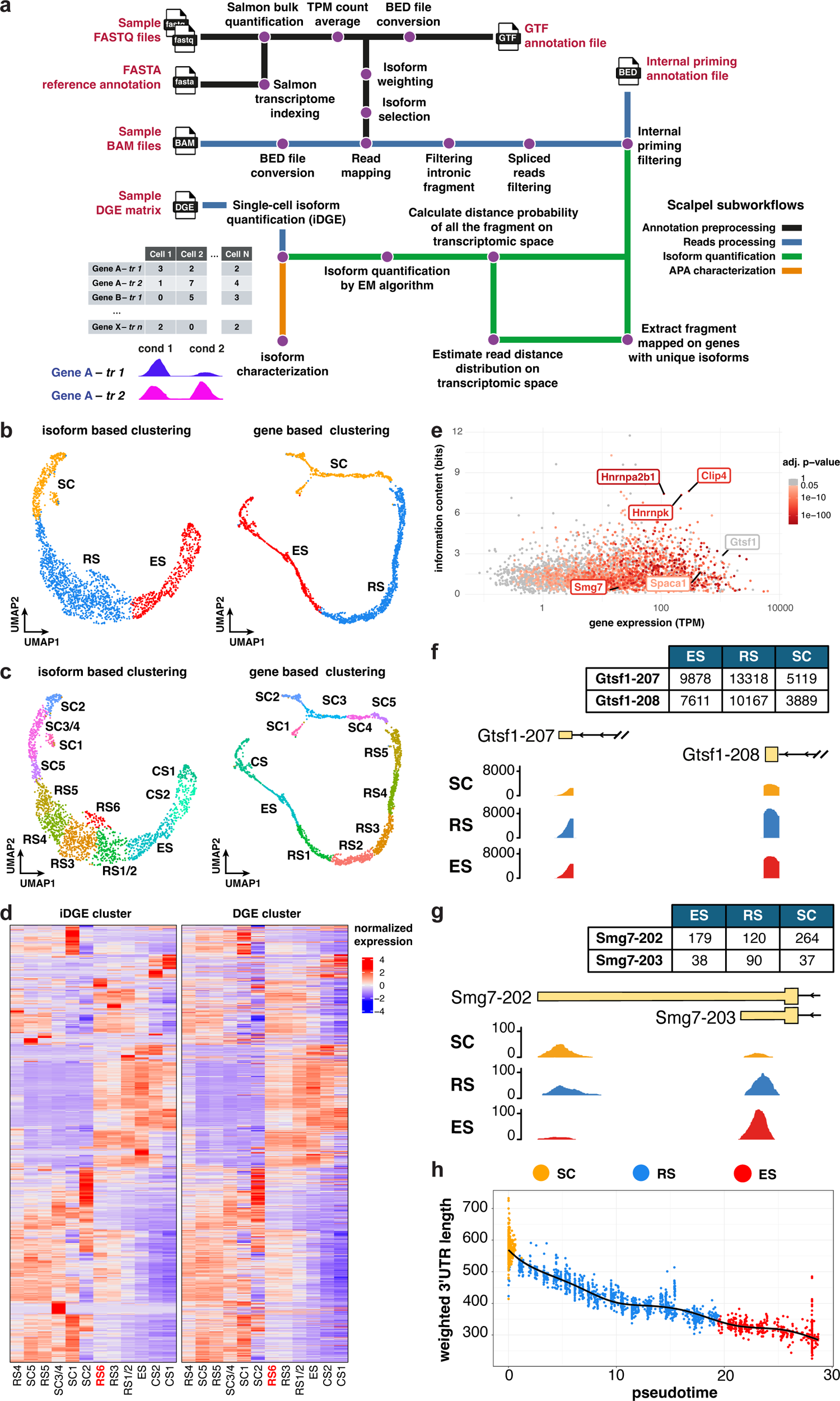
SCALPEL pipeline quantifies transcript isoforms at the single-cell level. **a.** SCALPEL Nextflow pipeline diagram. SCALPEL is composed of 4 workflows performing 1) annotation preprocessing (black line); 2) read preprocessing to discard artifacts and reads derived from pre-mRNAs (blue line); 3) quantification of isoforms in individual cells (green line); and 4) characterization of differential isoform usage (orange line). **b.** Cell types identified using isoform expression estimated by SCALPEL (left) or gene expression (right). Both analyses identify the same three cell populations: spermatocytes (SC), round spermatids (RS), and elongated spermatids (ES). **c.** High resolution clustering using isoform expression (left) identifies novel cell states (RS6) that cannot be identified using gene expression data (right). **d.** Heatmap showing the expression of cluster markers identified in the isoform based analysis (left) and their corresponding genes (right). The expression of some of the top RS6 isoform markers (black box, left) is not recapitulated with gene expression. Isoform expression distinguishes clusters more clearly than gene expression. **e.** Scatter plot showing the cell-type expression specificity relative to gene expression for all genes with two expressed isoforms. The higher the information content of a gene, the more cell-type specific its expression is. Colored dots represent genes whose isoform expression usage changes across conditions (Chi-squared test adjusted p-value < 0.05) **f, g.** SCALPEL quantification of isoform usage from *Gtsf1* **(f)** and Smg7 **(g)** genes. Coverage plots show the distribution of filtered reads along isoforms. *Gtsf1* **(f)** is an example of a gene with high cell type specificity identified in **e** whose isoforms show the same relative usage across cell types during mouse sperm cell differentiation. *Smg7* gene **(g)** has low cell type specificity as defined in **e** but significant changes in isoform usage across conditions. SCALPEL quantification and coverage plots show a gradual switch in isoform usage during the differentiation of SC to ES cells. **h.** Pseudotemporal ordering of cells confirms the overall shortening of 3’ UTRs during mouse sperm cell differentiation.

We used SCALPEL to investigate changes in isoform usage using a publicly available dataset on mouse sperm cell differentiation generated using 10x genomics platform^22^. In this dataset, SCALPEL identified 39,241 isoforms in 16,679 genes that were used for downstream analyses such as dimensionality reduction and clustering. The use of isoforms instead of genes for clustering analysis results in the identification of the same three main cell populations as using standard gene-based single-cell quantifications: elongated spermatids (ES), round spermatids (RS), and spermatocytes (SC) (Fig. 1b; 94% agreement in cell assignments). However, using a higher clustering resolution, we can identify new cell populations that cannot be identified using only gene expression-based approaches (Fig. 1c). We identified a new cell population of RS cells, RS6, that could not be distinguished using gene-based clustering (Fig. 1d, S2 a,b). GO term enrichment analysis using RS6 isoforms marker genes (Supplementary Table 1) identified biological processes associated to cilium organization and organelle assembly (Fig. S2c), which are essential processes for the differentiation and maturation of RS cells ^23,24^. DIU analysis across RS populations identified genes with differential isoform usage in RS6 cells, including changes in *Dnah3* and *Spaca1*, which are genes important for the morphological changes during sperm cell maturation such as the axonema and the flagellum (Fig. S2 d,e)^25,26^. Together, these results suggest that RS6 cell population corresponds to elongating spermatids^27^, an intermediate population state between RS and ES populations previously morphologically described in the literature that cannot be identified using single-cell gene expression profiles.

Given that isoform quantification is useful to identify new cell populations, we investigated how informative is isoform expression to define cell type identity. For this purpose, we computed for each gene its information content defined as the sum of the information from each of its isoforms^28^. The higher the information content of a gene, the more cell-type-specific the expression of its isoforms is. Using this approach, we noted that most genes had clear cell type specific expression bias (Fig. 1e). Manual inspection showed that in many cases all isoforms from the same gene showed similar expression changes across cell types, indicating that gene information content mainly reflects transcriptional changes (Fig. 1f). Thus, we used a chi-square test to identify genes in which the isoform usage changes across conditions, reflecting a regulation at the post-transcriptional level. Using this approach, we identified 4,214 genes displaying changes in isoform usage across cell types (Fig. 1e, red dots; Supplementary Table 2). One of these genes is *Smg7*, a gene that plays a crucial role in male germ cell differentiation in mice through its role in nonsense-mediated mRNA decay^25^. We observed a switch in isoform usage during cell differentiation where the long isoform of *Smg7* gene (Smg7-202) is progressively replaced by a shorter isoform (Smg7-203) (Fig. 1g).

Previous studies have shown that APA results in global 3’ UTR shortening during sperm cell differentiation^29,30^. Thus, we used SCALPEL predictions to assess if the observed changes in 3’UTR length reflected a coordinated shortening during sperm cell differentiation. We used the isoform quantification data to order cells according to pseudotime and measured the average 3’ UTR length of cells. In agreement with previous studies, we observed that overall, 3’ UTR length shortens while cells differentiate (Fig. 1h), indicating that SCALPEL predictions recapitulate known changes in 3’ UTR length during mouse spermatogenesis.

We next tested the performance of SCALPEL on a shallower dataset generated with Drop-seq platform. We profiled human induced pluripotent stem cells (iPSCs) and neural progenitor cells (NPCs) since it is known that APA changes significantly during neurogenesis^31^ and that miRNA regulation is very important in this process^32–34^. In this shallower dataset, SCALPEL quantified 59,796 isoforms in 18,617 genes and more than 10,000 genes with two or more isoforms. Isoform and gene-based analyses identified the same cell populations (Fig 2a). Differential isoform usage analysis identified 1,883 DIU genes with significant isoform expression changes between iPSCs and NPCs (Supplementary Table 3), highlighting that the drop in sequencing depth does not affect SCALPEL execution.

**Figure 2.**
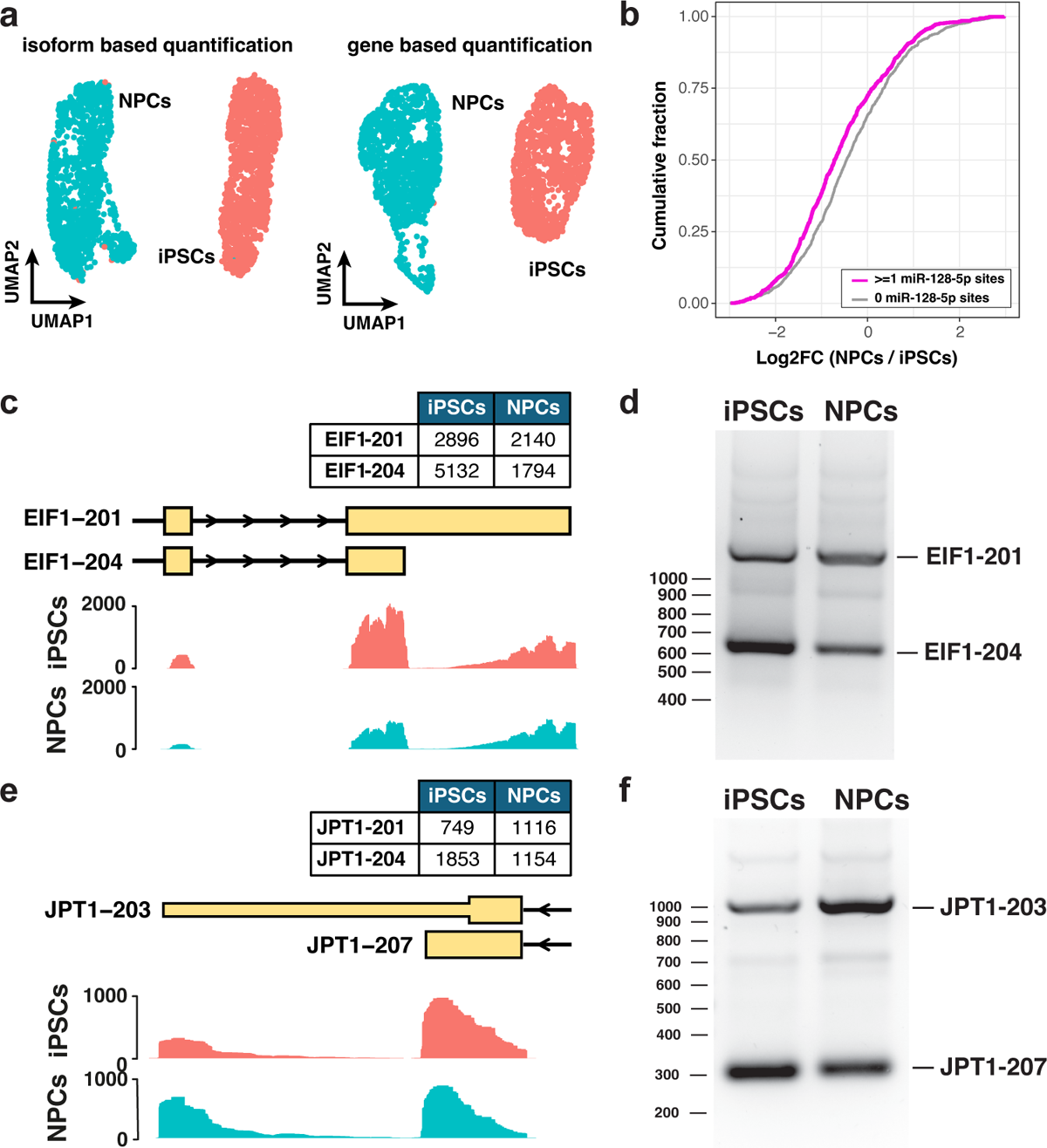
SCALPEL predictions on human Drop-seq data distinguish cell populations and provide insights into post-transcriptional regulatory mechanisms. **a.** UMAP plots based on isoform expression (left) and gene expression (right) similarly identify iPSCs (pink) and NPCs (blue). **b.** Cumulative distribution plot of log2fcs of isoforms containing miR-128-5p target sites (red) or without them (blue) from the same genes, indicating that changes in isoform usage can be attributed to miRNAs known to be implicated in neurogenesis. **c, e.** SCALPEL identifies a significant change in isoform usage in EIF1 **(c)** and JPT1 **(e)** genes between iPSCs and NPCs. Coverage plots show the distribution of filtered reads along isoforms. **d, f.** Relative isoform expression changes predicted by SCALPEL between iPSCs and NPCs in EIF1 and JPT1 genes can be experimentally validated in bulk using 3’RACE.

Given that isoforms with different 3’ends could contain different regulatory elements such miRNA target sites^2^, and that miRNAs usually downregulate their target RNAs^35^, we investigated if changes in isoform usage in NPCs compared to iPSCs was driven by miRNA function. To address this question, we downloaded the predicted miRNA target sites on the human genome from the MBS database^36^ and identified all isoforms targeted by miRNAs previously associated to NPCs^34^ (Supplementary Table 4). To investigate of miRNAs contribute to regulate isoform expression changes in NPCs, we compared the fold change distribution of isoforms containing miRNA target sites of miRNAs expressed in NPCs with those of non-targeted isoforms from the same genes. This analysis identified significant downregulation of isoforms targeted by let-7b, miR-124, miR-128, miR-199a, and miR-34 in NPCs compared to iPSCs (FDR < 0.05; Fig. 2b and Supplementary Table 5). This result suggests that miRNAs can explain some of the isoform expression changes predicted by SCALPEL during the differentiation of iPSCs to NPSc.

Considering that these two cell types can be easily distinguished experimentally, we decided to use this dataset to experimentally validate SCALPEL predictions. To this end, we selected six genes predicted to have changes in isoform usage by SCALPEL across cell types and validated the isoforms expressed by these genes in NPCs and iPSCs using 3’RACE. This analysis validated the changes in isoform usage in two of the six genes (Fig. 2c-f) and detected all isoforms predicted by SCALPEL in two of the remaining four genes (Fig. S3).

Finally, we benchmarked SCALPEL performance against existing tools developed to quantify APA in scRNA-seq data^20,37–41^. Considering the underlying quantification strategy, these methods can be divided into peak-calling based tools (Sierra, scAPA, scAPAtrap, SCAPTURE, and scDapars), and isoform quantification tools (scUTRquant) (Supplementary Table 6). Following the quantification of peaks or isoforms according to the default parameters of each tool, we performed DIU analysis across pairs of cell types (Fig. S4a). In the case of scUTRquant^20^, which uses an extended curated 3’ UTR annotation (3’ UTRome), we performed the benchmarking using both the 3’ UTRome (scUTRquant) and the same annotation as the other tools (scUTRquant*). Overall, our analyses show a clear difference in sensitivity between peak and isoform-based methods (Fig. 3a). Peak-based methods quantified fewer genes and isoforms than isoform-based methods, which all showed similar sensitivities. Most of the sensitivity differences between peak and isoform tools can be explained by the constraints of the prediction methods. For instance, the low number of isoforms quantified by scDapars^42^ and scAPA^19^ can be explained because these tools only predict PAS in annotated 3’ UTRs. In contrast, the large number of peaks detected by scAPAtrap^17^ is explained because its predictions are not restricted to the gene annotations (Supplementary Tables 7,8).

**Figure 3.**
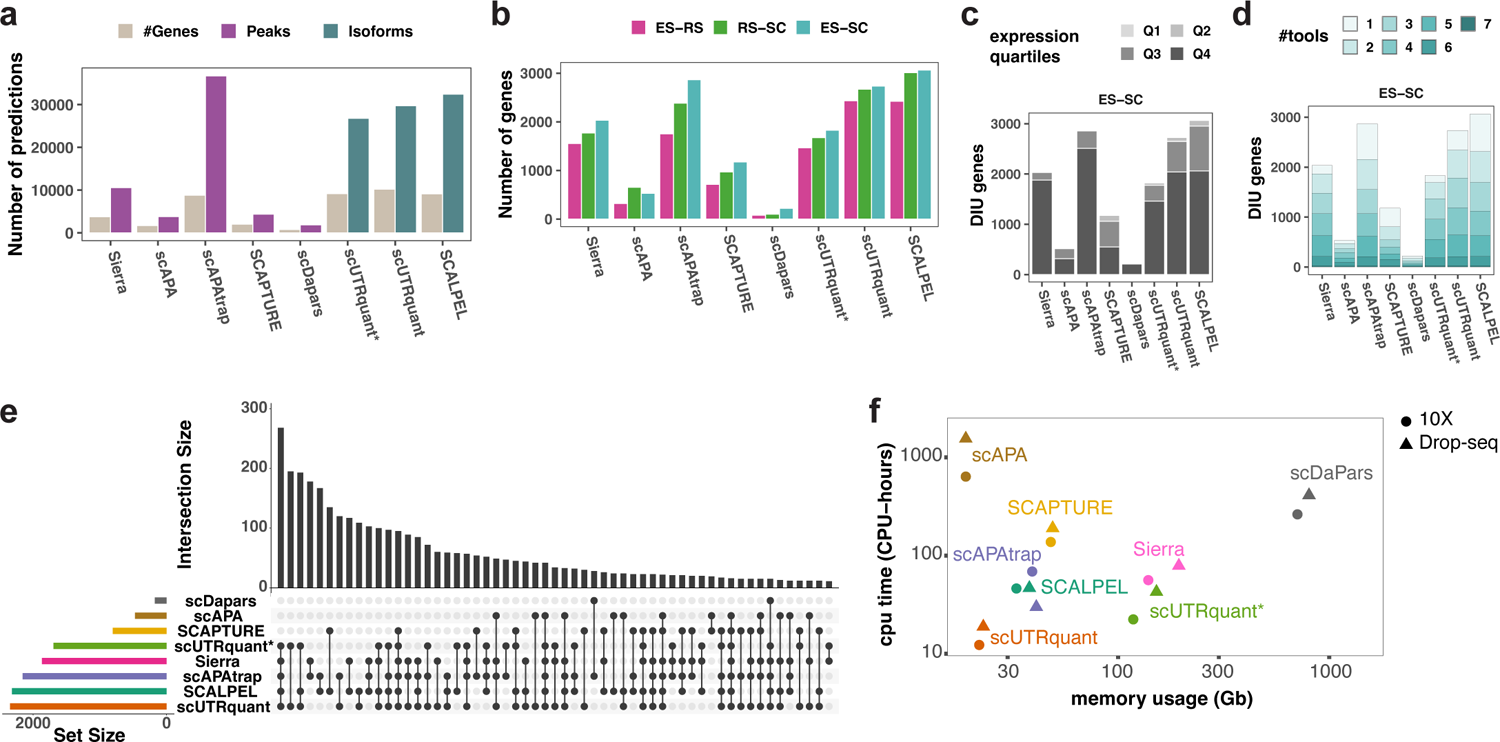
SCALPEL benchmark against existing tools for APA and isoform quantification at the single-cell level. **a.** Numbers of genes (grey), peaks (purple) and isoforms (sea green) quantified by each tool in the 10x mouse spermatogenesis dataset. scUTRquant has been run using the UTRome annotation (scUTRquant) and the same annotation as all other tools (scUTRquant*). **b.** Number of DIU genes between ES and RS cells (dark pink), RS and SC cells (green) and ES and SC cells (light blue). In all comparisons SCALPEL predicts more DIU genes than all other tools. **c.** Distribution of DIU genes between ES and SC according to their raw UMI counts split by quantiles (Q1: 4-37, Q2: 37-277, Q3: 277-1618, Q4: 1618-423964). Predicted DIU genes by all tools are mainly among highly expressed genes (Q3 and Q4 quartiles). **d.** Agreement in the identification of DIU genes between ES and SC clusters across all tools. 75% of DIU genes predicted by SCALPEL are also predicted by one or more of other tools. **e.** UpSet plot showing the agreement in the number of predicted DIU genes between ES and SC across all tools. All intersection sets including more than 10 genes are shown. **f.** Comparison of the CPU time vs the memory usage for the different tools. Except for scUTRquant, which uses an already processed annotation, SCALPEL is the most efficient tool of all the ones tested (less memory and CPU time used).

Next, we identified DIU genes for each pair of cell types. SCALPEL identified the highest number of DIU genes, closely followed by scUTRquant and scAPAtrap (Fig. 3b). This higher sensitivity is not driven by the quantification of lowly expressed genes, as most DIU genes are highly expressed (top 50%; Fig. 3c and S4b,e). Here, it is important to note that the number of DIU genes detected by scUTRquant using the standard gene annotation (scUTRquant*) is clearly reduced, indicating that the higher sensitivity of scUTRquant can be directly attributed to the use of an extended annotation (3’ UTRome) and not to the algorithm *per se*. Finally, we investigated the agreement in the prediction of genes with differential peak or isoform usage across tools. We observed a substantial overlap in the predictions of SCALPEL with other tools, with more than 70% of SCALPEL predictions supported by one or more tools (Fig. 3d,e; Fig. S4b,e). SCALPEL and scUTRquant showed the highest agreement on the identified DIU genes across the cell types (Fig. S4c,f,j-l). When performing the benchmark on the neuronal dataset, SCALPEL and scAPAtrap showed higher sensitivity in the identification of DIU genes while keeping a high degree of agreement with other tools (Fig. S4g-i, m). In this case, although scUTRquant quantified a high number of isoforms (40,002) and genes with multiple isoforms (10,481), only 246 DIU genes were identified between NPCs and iPSCs (Fig. S4 g-i). This drop in the number of detected DIU genes is likely arising from stringent default parameters which discard genes expressed in a few cells (*minCellsPerGene*=50).

Finally, we decided to compare the performance of SCALPEL in terms of execution requirements and runtime. SCALPEL run time and memory usage is comparable or better than most of the tools analyzed, and only scUTRquant is faster and more memory efficient than SCALPEL. Yet, this can be directly linked to the presence of an already processed annotation as both memory and execution time increase when no 3’ UTRome is provided to scUTRquant (scUTRquant*, Fig. 3f, Supplementary Tables 9, 10).

## Conclusions

In this manuscript we have presented SCALPEL, a new tool for sensitive isoform quantification and visualization using conventional 3’ based scRNA-seq data. Comparison of SCALPEL to other existing tools shows that SCALPEL quantifies more isoforms and detects more genes with changes in isoform usage (Fig. 3a). While we do not have a ground truth dataset to assess the accuracy of the predictions, we show that SCALPEL predictions have a high agreement with that of the other tools (Fig. 3d,e and Fig. S4), which has commonly been used as a measure of accuracy in these cases, and that SCALPEL’s predictions can be validated experimentally (Fig. 2 d, f, Fig. S3).

We demonstrate that the iDGE provided by SCALPEL can be used to perform standard single-cell analysis such as dimensionality reduction, clustering and pseudotime analysis (Fig. 1 b,c,h and Fig. 2a). We also show that isoform quantification can be used to gain new insights about cell populations and identify, for instance, novel cell states that cannot distinguished using standard single-cell gene quantification data. (Fig. 1c and Fig S2). Furthermore, we highlight how SCALPEL predictions can be used to investigate mechanisms of gene regulation at the single-cell level. SCALPEL predictions recapitulate known changes in 3’ UTR length during cell differentiation (Fig. 1h) and reflect miRNA function at the single-cell level, as changes in isoform usage across cell types can by directly linked to the presence of cell type specific miRNAs (Fig. 2b). Together, our work highlights how SCALPEL expands the current scRNA-seq toolset to explore post-transcriptional gene regulation in individual cells from different species, tissues, and technologies to advance our knowledge on gene regulation from the bulk to the single-cell level.

## Methods

### Annotation preprocessing

3’ based scRNA-seq protocols use oligo(dT)s to capture polyadenylated RNAs, which introduces a bias in the location of reads towards the 3’ end of the RNAs (Fig. S5). Thus, we truncated the annotated isoforms in the existing annotation and restricted the quantification to isoforms that are different at the 3’ end. Using GENCODE^43^ annotation as reference, we truncated all isoforms to include the last 600 nucleotides of spliced sequence from their 3’ end, which is the region that displays coverage by the 3’ tag-based scRNA-seq data (Fig. S5). Then, we collapsed truncated isoforms with exact intron/exon boundaries and fewer than 30 nucleotides differences in their 3’ end coordinates. When multiple isoforms were collapsed, we kept the name of the isoform having the higher expression according to pseudobulk quantification. For this purpose, we used *salmon quant* v0.14^44^ with default parameters to quantify all isoforms in bulk using as input the scRNA-seq bam files provided. For the analysis of multiple samples, isoform expression was averaged across all samples.

### Read preprocessing

We processed all the input BAM files containing aligned and tagged scRNA-seq reads to discard artifacts and reads not supporting annotated transcripts. First, we used *samtools* v1.19.2^45^ to split the input BAM files by chromosome using the command *samtools view* and converted them into BED files using *bam2bed* command from BEDOPS v2.4.41^46^. We included all reads in the bed file (option –all-reads) and split them into separate entries (option –split) if contained Ns in the CIGAR line (*i.e.* spliced reads). We used the function *findOverlaps* from GenomicRanges R package v1.50.0^47^ to overlap the reads with the set of selected isoforms using default parameters. Given that all reads with the same cell barcode (BC) and unique molecular identifier (UMI) likely were generated from the same original RNA molecule, we grouped them into a unique fragment and jointly evaluated them during isoform quantification (Fig. S1). We defined the genomic coordinates of each fragment as the most 5’ and 3’ coordinates of its associated reads. We discarded all the fragments overlapping intronic and intergenic regions except those extending the 3’ end of the gene, and spliced reads not supporting annotated exon-exon junctions, as they were considered to come from pre-mRNAs or unannotated transcripts. To avoid biases in the quantification of the isoforms due to reads mapping to IP locations in the genome, we discarded all the fragments that could arise from these sites. For that purpose, we scanned the whole genome using a custom Perl script and identified putative IP locations as regions containing six or more consecutive adenosines in a window of 10 nucleotides. We discarded all the fragments located upstream from an IP site. In this case, we only considered IP sites located more than 60 nucleotides upstream of an annotated 3’ end in the transcriptomic space.

### Quantification of isoforms at the single-cell level

First, we used all genes with only one detected isoform to assess the empiric distance distribution of scRNA-seq reads with respect to annotated isoform 3’ ends. Considering defined intervals of 30 nt, we calculated the distribution of read 3’ends relative to annotated 3’ends by dividing the number of distinct 3’ends in each bin in the transcriptomic space divided by the total number of 3’ends. Using this probability distribution, we assigned to each read a probability to come from a specific isoform based on its relative distance to the 3’ end. Considering that each fragment is composed of one or more reads, we defined the probability of a fragment *f* to belong to an isoform *k* for a gene *g* as:

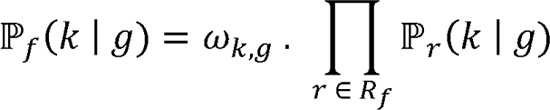

where *R*_*f*_ is the set of reads associated with the fragment *f*, ℙ_*r*_(*k* | *g*) is the probability that read *r* is associated to the isoform *k*, and ω_*k*,*g*_ is the weight associated to the isoform *k* and gene *g* based on the pseudobulk quantification.

We computed the weight ω_*k*,*g*_ of an isoform *k* and gene *g* as

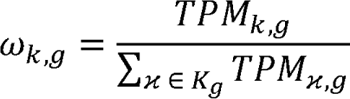

where *K*_*g*_ is the set of isoforms for the gene g and *TPM*_*k,g*_ is the transcript per million counts of isoform *k* and gene *g* according to Salmon^44^.

Hence, the probability of a set of fragments *F*_*g,c*_ for gene *g* and cell *c* given a fixed set of isoforms *K*_*g*_ and associated relative abundances values (θ) is given by:

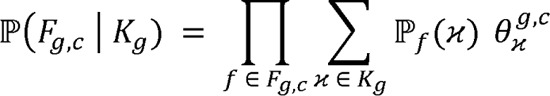

where *F*_*g,c*_ is a set of fragments for a gene *g* and cell *c*, ℙ_*f*_(x) represents the probability of a fragment *f* belongs to isoform x, and θ ^*g,c*^_x_ the isoform x relative expression for a gene *g* and cell *c*.

Consequently, we estimated the isoform relative expression values (θ) by maximizing the log-likelihood function using a standard Expectation Maximization (EM) algorithm. The maximum likelihood estimators ^θ^^ for the isoform relative abundance values are given by:

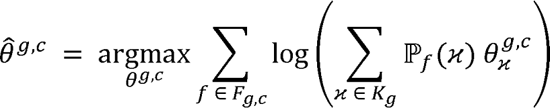

For each gene g and cell c, the EM algorithm proceeds as follows:

1. Initialization of value θ_*k*_^*g,c*^ for each isoform *k* of the gene g as 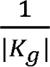
2. Iterate until convergence:

a. E-step: Calculation of posterior probability for each fragment *f* ∈ *F*_*g,c*_ to belong to the isoform *k* given that it comes from a gene *g* and a cell *c* as

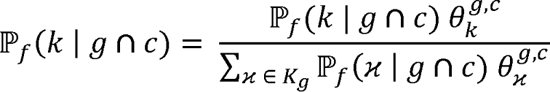
b. M-step: Estimate the isoform relative expressions θ^*g,c*^

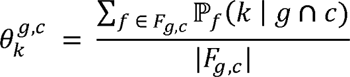

The convergence state of the EM algorithm was settled by a stop criterion condition. This condition was reached when the maximum difference of the estimated relative abundance between two iterations was equal to 0.001. All the isoforms with a null weight value were discarded from the annotation set. To generate isoform expression values for the iDGE per gene the estimated isoform probabilities were multiplied by the UMI counts assigned to the into the original DGE (Fig. S1).

### Isoform entropy and gene information content calculation

In order to deconvolute the DGE into an iDGE, SCALPEL estimates the isoform relative expression distributions for each gene and cell (θ, see above). Considering a fragment *f*^-^ across the set of all the fragments F, we can define the conditional probability of fragment *f*^-^ to originate from an isoform *k* given it comes from a gene *g* and cell *c* as:

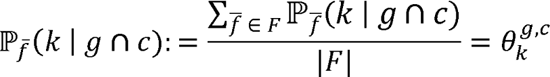

where θ ^*g,c*^ is the isoform *k* relative expression estimated by SCALPEL for gene *g* in cell *c*. Taking this into account, the probability of fragment *f*^-^ to originate from isoform *k* of gene *g* given that it originates from cell *c* can be derived as

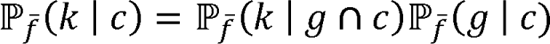

where the probability of fragment *f*^-^ to originate from any isoform of gene *g* given it originates from cell *c* can be calculated from the iDGE as:

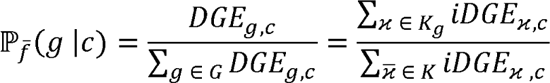

Where *DGE*_*g,c*_ denotes the number of UMIs assigned to gene *g* in cell c, *G* the set of all the genes, *iDGE*_x,*c*_ the number of UMIs assigned to isoform x in cell c, *K* the set of all the isoforms across all the genes.

Applying Bayes’ theorem, the probability of fragment *f*^-^ to originate from cell *c* given it originates from isoform *k* can be derived as:

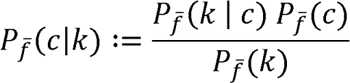

where the probabilities *Pf*^-^(*c*) of fragment *f* to originate from cell *c* and *Pf*^-^(*k*) of fragment *f* to originate from isoform *k* can be estimated from the iDGE corresponding column and row sum fractions, respectively.

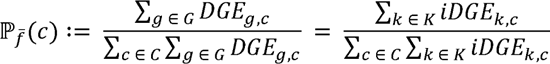

where *C* is the set of all the cells, and

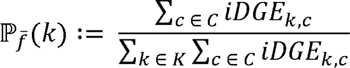

Given the cell to cell cluster mapping ϕ:C↦L, where *L* is the set of all cell clusters, the probability of fragment *f*^-^ to originate from a cell in cluster *c*, given it originates from isoform *k* can be derived by summing up the corresponding cell probabilities:

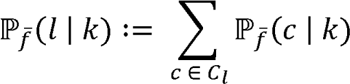

where *C*_*c*_ is the set of the cells associated to the cluster *c*.

Based on those conditional probabilities, each isoform’s entropy across cell cluster is defined as:

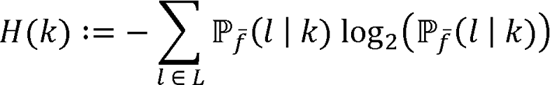

If all fragments originating from a given isoform originate from cells of the same cell type, the corresponding entropy is minimal, at a value of

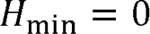

bits. If an isoform *k* is expressed perfectly equal across all clusters, the maximum isoform entropy

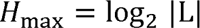

is reached, where |L| denotes the number of cells clusters.

The isoform-level entropies were summarized to the gene level to quantify the randomness of a gene’s isoform distribution across cell types:

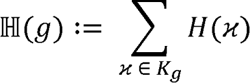

While the minimal gene entropy

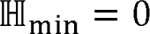

is gene-independent, its upper bound depends on the number of expressed isoforms |*K*_*g*_| and thus differs across genes:

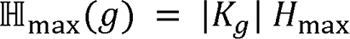

The difference from that maximum entropy defines a gene’s information content across isoforms and cell types/clusters

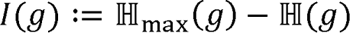

### Detection of differential isoform usage between cell clusters

For the identification of differential isoform expression between clusters of cells, we implemented the function *FindIsoforms*. This function selects all genes with at least two isoforms expressed and performs a Chi-squared test to assess if the read distribution across isoforms in the same between clusters. In these analyses, all isoforms representing less than 10% of the expression of a gene in at least one condition were discarded. We selected significant DIU genes with a false discovery rate (FDR) < 0.05.

### Average 3’ UTR length calculation

We extracted from the iDGE all protein-coding isoforms and computed the lengths of the corresponding 3’ UTR regions from the reference annotation. For each gene *g*, we calculated the gene average 3’ UTR length τ_*g*_ in a cell *c* as

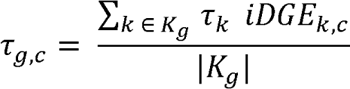

where τ_*k*_ is the 3’ UTR length of isoform *k*, *iDGE*_*k,c*_are the UMI counts of isoform *k*.

Next, we calculated the average weighted 3’ UTR isoform length within each cell τ_*c*_ for all the expressed isoforms as

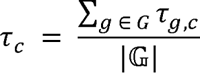

where 𝔾 is the set of all genes with protein-coding isoforms expressed.

### Generation of read coverage plots

SCALPEL outputs a BAM file including the reads used for isoform quantification. Within SCALPEL framework, we have implemented the function *CoveragePlot* in R to visualize the read coverage on the isoforms using a transcriptome annotation in GTF format and BAM files. We generated the visualization tracks using the R Gviz library v1.46.0^48^.

### Analysis of mouse spermatogenesis 10x scRNA-seq data

We downloaded the 10x scRNA-seq samples from male mouse germline^22^ from GEO database (accession number GSE104556) and processed them using Cell Ranger v7.1.0^49^, using mm10 mouse genome assembly^50^ and GENCODE M21^43^ as reference annotation. We merged the processed data and analyzed them jointly using Seurat v5.0.0^51^. We restricted the analysis to the set of annotated cells from a previously study^19^. The final Seurat object contained 2,042 cells and 22,433 genes. We performed dimensionality reduction on the DGE using 2,000 genes as variable features and used the first 9 principal components (PCs) to build the kNN graph and compute a UMAP. We used the function *FindClusters* with a resolution of 0.03 to identify 3 clusters. We compared our clustering results with the cell clustering from the previous study on the same data by calculating a Jaccard Index score between the cell barcode tags clusters and annotated the defined cell clusters in our analysis as ES, RS and SC according to their previous corresponding annotation. Next, we filtered the input BAM file to retain the 2,042 annotated cells included in the previous analyses using *samtools view - D CB* from samtools v1.19.2^45^ and extracted the DGE matrix for SCALPEL analysis. Following the execution of SCALPEL, we generated a Seurat object containing the same 2,042 cells, 19,797 genes and 70,793 isoforms. For downstream analyses, we discarded all isoforms expressed in less than four cells and cells with less than three isoforms expressed using *CreateSeuratObject* function (option min.cells=4, min.features=3). The final Seurat object included 40,750 isoforms of 16,752 genes. We performed dimensionality reduction on the iDGE using 2,000 isoforms as variable features, and we used 11 PCs for the generation of the kNN graph for the UMAP. We used a resolution of 0.05 to perform the clustering and identified three cell populations. We used the approach described above to annotate the cell clusters. We compared the iDGE clusters with the DGE clusters using a Jaccard index.

We increased the clustering resolution in the gene and the isoform-based analyses to 0.4 and 0.9 respectively to identify new cell states. Using the markers from Lukassen et al. ^22^, we annotated the cell clusters of the gene-based analysis. For the isoform analysis, we calculated a Jaccard index score between all gene and isoform clusters (Fig. S2) and assigned to each isoform-based cluster the identity of the most similar cluster. We performed a differential isoforms analysis using *FindAllMarkers* function (option min.pct=3, p_val_adj<0.05) on the iDGE to identify isoforms markers for each cluster. We visualized the top 100 isoform markers average expression within each cluster in the iDGE using the *Heatmap* function from the ComplexHeatmap^52^ package along the average expression of their corresponding genes in the DGE clusters (Fig 1d).

### Differentiation and characterization of iPSCs and NPCs using Drop-seq technology

We differentiated human iPSCs towards NPCs using a protocol previously established in the lab^53,54^. Briefly, we differentiated iPSCs to neuroepithelial cells over a period of ten days by dual SMAD inhibition using neural maintenance medium (1:1 ratio of DMEM/F-12 GlutaMAX (Gibco, #10565018) and neurobasal (Gibco, #21103049) medium complemented with 0.5x N-2 (Gibco, #17502048), 0.5x B-27 (Gibco, #17504044), 2.5 μg/ml insulin (Sigma, #I9278), 100 mM L-glutamine (Gibco, #35050061), 50 μM non-essential amino acids (Lonza, #BE13-114E), 50 μM 2-mercaptoethanol (Gibco, #31350010), 50 U/ml penicillin and 50 mg/ml streptomycin (Gibco, #15140122)) supplemented with 500 ng/ml noggin (R&D Systems, # 3344-NG-050), 1 μM Dorsomorphin (StemCell technologies, #72102) and 10 μM SB431542 (Calbiochem, # 616461). After the initial neural induction step, we differentiated the cells to NPCs by neural maintenance medium replacement up to day 22. We cryopreserved iPSCs and NPCs in fetal bovine serum or neural maintenance medium supplemented with 10% of DMSO for iPSC or NPCs respectively. Before single-cell encapsulation using NADIA instrument (Dolomite Bio), we thawed, filtered and counted the samples. We retrotranscribed RNAs captured with the oligo(dT) for cDNA library preparation. Finally, we sequenced the final Illumina tagged libraries on an Illumina NextSeq 550 sequencer using the NextSeq 550 High Output v2 Kit (75 cycles) (Illumina, #20024906) in pared-end mode; read 1 of 20 bp with custom primer Read1CustSeqB^55^, read 2 of 64 bp and 8 bp for i7 index.

### scRNA-seq analysis of Drop-seq data

We processed the iPSC and NPC scRNA-seq libraries using Drop-seq tools v2.5.1^55^ pipeline to generate DGE matrices. We merged the FASTQ files containing paired end reads into a single unaligned BAM file using Picard tools *v2.*27.4^56^. We tagged the reads with cell and the molecular barcodes, trimmed them at the 5’ end to remove adapter sequences and at the 3’ end to remove polyA tails, and mapped them to the human genome (*GRCh38*) with STAR *v2.7.10*^57^. We tagged the resulting BAM files with the annotation metadata using the human GENCODE v41^58^ annotation as reference. Finally, we performed the cell barcode correction using the programs *DetectBeadSubstitutionError* and *DetectBeadSynthesisErrors* with default parameters. To estimate the number of cells obtained, we used a knee plot considering the top 3,000 cell barcodes and generated a DGE count matrix for each sample.

We used Seurat v5.0.0^51^ and R 4.3.2^59^ to merge the DGEs and preprocess the scRNA-seq data. We discarded all genes expressed in less than four cells and all cells with less than three genes expressed. We also discarded low quality cells with less than 300 UMIs, less than 300 genes and more than 5% mitochondrial gene, and cell artifacts with more than 20,000 UMIs and 7,000 genes. The final Seurat object contained 2,535 cells and 19,103 genes. We performed a dimensionality reduction analysis using the 2,000 most variable genes to calculate 50 PCs. We used the *ElbowPlot* function to manually inspect the amount of variability explained by each PC and selected the first 9 PCs to build the kNN graph and compute the UMAP plot.

### Trajectory inference analysis

We used the iDGE-based Seurat object to derive a pseudo temporal ordering of the cells using Monocle3 v1.3.4^60^. First, we converted the Seurat object into a CellDataSet object which includes the cluster annotation and the UMAP embedding previously computed. Next, we fit the principal trajectory graph within each cluster partition using the function *learn_graph*. Finally, we calculated the pseudotime values using the function *pseudotime* and the ES cells as root cells.

### GO term enrichment analysis

We performed GO term enrichment analysis using the R package enrichR v3.2^61^. The reference used for the enrichment analysis was the GO_Biological_Process_2023^62^. We selected all the GO terms with an adjusted p-value inferior to 0.05 and visualized the associated adjusted p-value and enrichR combined scores using ggplot2^63^.

### Identification of miRNA signatures in differentially regulated isoforms

To obtain isoform-level identification of miRNA target sites, we overlapped the genome-wide miRNA target site annotation included in the MBS database^36^ with the GENCODE annotation reference v41^58^ (hg38). Target sites of different miRNAs were grouped by their seed sequences according to miRBase v22.1^64^ The seed-target isoform pairs were used for downstream analyses. Using a set of neurogenesis-related miRNA^34^, we filtered genes displaying changes in isoform usage between iPSCs and NPCs as predicted by SCALPEL with at least one isoform targeted by a neurogenesis-related miRNA and one non-targeted isoform. We normalized the isoform abundances by the number of cells in NPCs and iPSCs and used these values to calculate their log2 fold changes (log2fcs). For each miRNA, we used a Kolmogorov-Smirnov test to check for differences in the cumulative distribution of log2fcs between targeted and non-targeted isoforms from the same set of genes. The resulting p-values were FDR-adjusted using Benjamini-Hochberg correction.

### Isoform validation using nested PCRs

To validate the changes of isoform usage between the iPSCs and NPCs predicted by SCALPEL, we performed nested PCR as previously described^65,66^. One µg of total mRNA extracted using Maxwell® RSC simplyRNA Cells kit protocol (Promega Corporation, #AS1340) was used as input RNA for the cDNA synthesis using an oligo(dT)-adaptor sequence TAP-VN as a primer for the reverse transcription. For the first nested PCR, one µL of 1:10 cDNA dilution was used, with a gene-specific primer (GSP) which is shared by all isoforms and an adaptor primer (AP) as a reverse primer. The second nested PCR was performed with one µL of a 1:10 dilution of the first PCR using a second gene-specific primer and a second adaptor primer (MAP) as a reverse primer. This second nested PCR reaction that anneals 3’ to the first GSP is essential to reduce the amplification of undesired products^67^. The resulting nested PCR products are resolved by an agarose gel. Primer sequences are provided in Supplementary Table 11.

### Benchmark analysis

We downloaded each of the benchmarked tools from their respective GitHub repository. For each tool, all commands for its default execution were integrated into Nextflow workflows and were executed using the default parameters indicated by the authors. We performed the benchmark analysis on the preprocessed mouse spermatogenesis scRNA-seq^22^ data using the GENCODE vM21^58^ GTF and transcriptome FASTA files as reference annotation. Additionally, we performed a second benchmark analysis on the preprocessed neuronal cell differentiation scRNA-seq Drop-seq data using the analogous reference files obtained from GENCODE v41. As scAPA only considers disjoint 3’UTR regions annotation from the hg19 version of the human genome for the annotation of its quantified peaks, we generated a new annotation of non-overlapping 3’UTR regions using GENCODE v41 and *GenomicRanges* R package. We intersected all the peaks detected by scAPA^19^ following its peak calling process to this new reference annotation. Following Dapars2 v2.1^68^ default procedure, we downloaded the gene region annotation reference for the human and mouse genome (GRCh38, mm10) using the UCSC Table browser. Then, we extracted 3’ UTR regions from the gene annotation using Dapars2 script *DaPars_Extract_Anno*. Finally, we calculated the raw percentage of distal PAS sites usage index (PDUI) values using Dapars2 script *DaPars2_Multi_Sample_Multi_Chr* and provided them to scDapars^42^ to infers their expression at single-cell level. scUTRquant^20^ was executed using the target transcriptome annotation files provided in the GitHub repository for the human and mouse genomes (hg38 and mm10). These files included high-confidence cleavage sites called from the Human Cell Landscape and Mouse Cell Atlas dataset^20^. Additionally, we reran scUTRquant^19^ using a custom target transcriptomic annotation generated from the input genome annotation using the Bioconductor package *txcutr*^69^ (v1.8.0). For each tool, we performed a differential peak or isoform usage analysis with default parameters for all genes with at least two peaks or isoforms detected. As Sierra did not allow for the analysis of differential isoform usage across multiple cell types, we performed pairwise comparisons across the three cell types (ES-RS, RS-SC and ES-SC). Next, we compared the DIU genes between the iPSCs and NPCs for the neurogenesis Drop-seq dataset. We discarded all the DIU genes with an adjusted p-value greater than 10%. For each comparison test, we generated the UpSet plots using the R library UpSetR v1.4.0^70^ for the set of DIU genes co-detected by at least two tools. We applied a cutoff threshold of 10 genes for each intersection set.

### Software used

SCALPEL is implemented in Python and R^59^. We performed all the statistical tests in this manuscript using R v4.3.2^59^.

## Code availability

The code of SCALPEL and the pipeline to benchmark all tools presented in this manuscript are available on GitHub (https://github.com/p-CMRC-LAB/SCALPEL).

## Data availability

The Drop-seq data from iPSCs and NPCs generated within this project are available on GEO under accession number GSE268222.

## Author contributions

FA developed SCALPEL. FA, MS, AJG, LL and MP designed the computational analyses. FA, MS and AJG performed computational analyses. FA, MS, AJG and MP interpreted the data. AGF generated scRNA-seq data. SMFM, FA and MP selected candidate genes for validation. SMFM performed experimental validations. MP designed the project, acquired funding, supervised, and coordinated the work. FA and MP wrote the initial draft with input from all authors. All authors reviewed the final manuscript.

## Acknowledgements

The authors thank all members from the Plass Lab for useful comments and critical discussions. We thank Loris Mularoni from the Regenerative Medicine Program for his support with the cluster management. We also thank Nicole Grieger for her help setting up 3’RACE and Yvonne Richaud-Patin and Dr. Zomeño from the Regenerative Medicine Program and IDIBELL’s Advanced Cell and Tissue Culture platform respectively for their help with iPSC cell culture. This research was funded by research projects from the State R&D Program Research Challenges from the Spanish Ministry of Science, Innovation and Universities (Project: PID2019-108580RA-I00 funded by MICIU/AEI /10.13039/501100011033 and Project: PID2022-139580OB-I00 funded by MICIU/AEI /10.13039/501100011033 and FEDER and EU) M.P. work is supported by a Ramón y Cajal contract of the Spanish Ministry of Science, Innovation and Universities (Grant: RYC2018-024564-I funded by MICIU/AEI/10.13039/501100011033 and by “El FSE invierte en tu futuro”). F. A. work is supported by a predoctoral contract of the Spanish Ministry of Science, Innovation and Universities (Grant: PRE2020-094049 funded by MICIU/AEI /10.13039/501100011033 and by “FSE invierte en tu futuro”). We thank CERCA Program/Generalitat de Catalunya for IDIBELL institutional support.

