## Supplementary Figures for "Quantification of transcript isoforms at the single-cell level using SCALPEL"

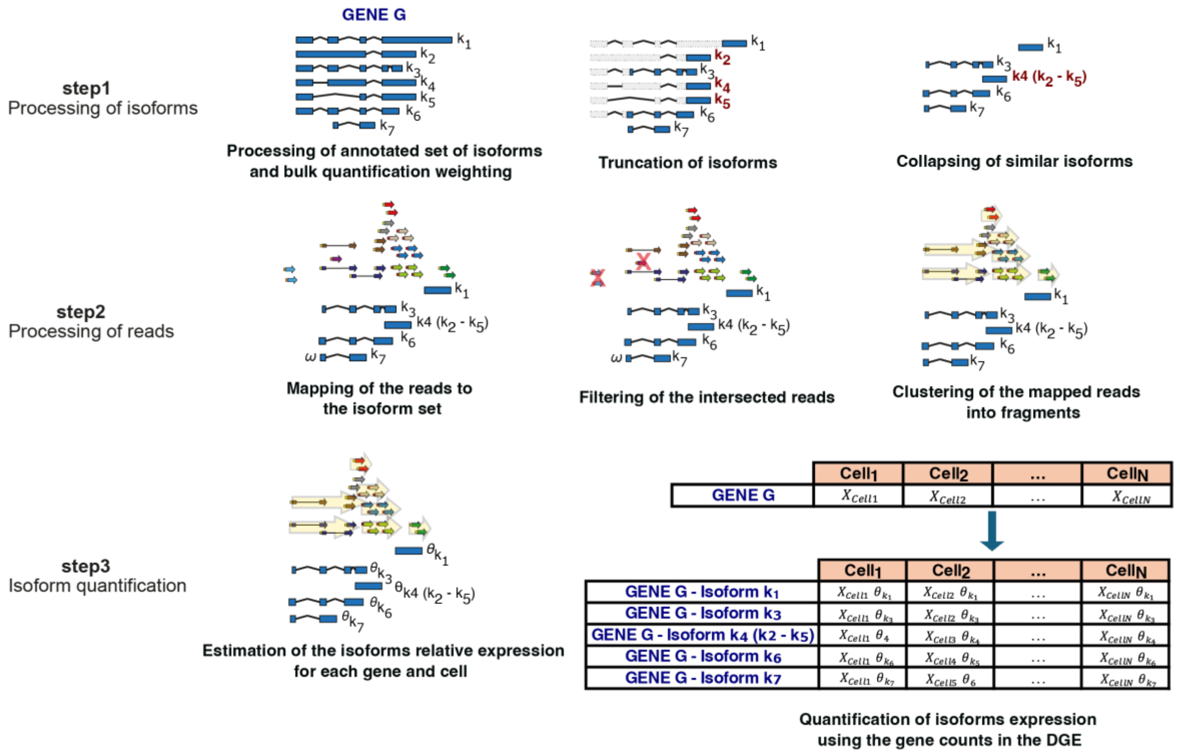


**Figure S1. Detailed SCALPEL workflow.** SCALPEL performs isoform quantification in 3 steps. Step 1. Isoforms are truncated to the last 600 nt before collapsing similar isoforms. Step 2. All the reads annotated to genes in the BAM file are mapped to the set of collapsed isoforms. Intronic or IP-derived reads are discarded. Reads with the same cell barcode and UMI are then assembled in fragments. 3. Relative isoform abundances are estimated for each cell using the fragment probabilities. Once isoform probabilities are estimated, final isoform expression is calculated by multiplying the relative isoform abundances by the UMI counts assigned to each gene in each cell.

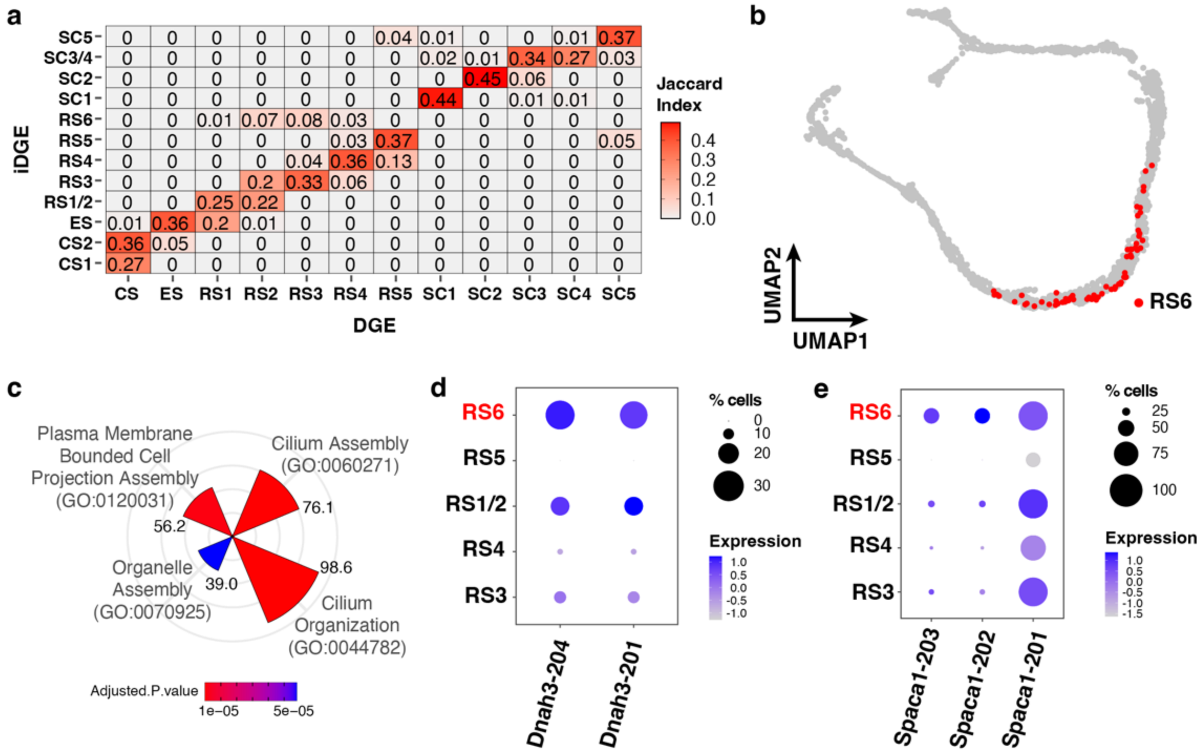


**Figure S2. Differences between gene- and isoform-based clustering solutions. a.** Similarity in the cell classification across different clusters between the isoform- (iDGE) and the gene- (DGE) based analyses. All the clusters in the iDGE analysis have a one to one correspondence with DGE clusters. **b.** Cells from the RS6 cluster are highlighted on the UMAP generated using gene expression data. In this plot, RS6 cells are spread across multiple RS clusters (RS1-RS4) and are not located within any specific cluster. **c.** GO term enrichment analysis using RS6 marker isoforms identified significant terms associated to cilium organization and cell differentiation. In the circular barplot, GO terms are arranged according to their combined enrichR score and colored according to their adjusted p-value. **d,e.** DotPlot of RS6 marker isoforms of Dnah3 **(d)** and Spaca1 **(e)** genes across RS cell populations highlights differences in the expression of these isoforms across RS cell populations.

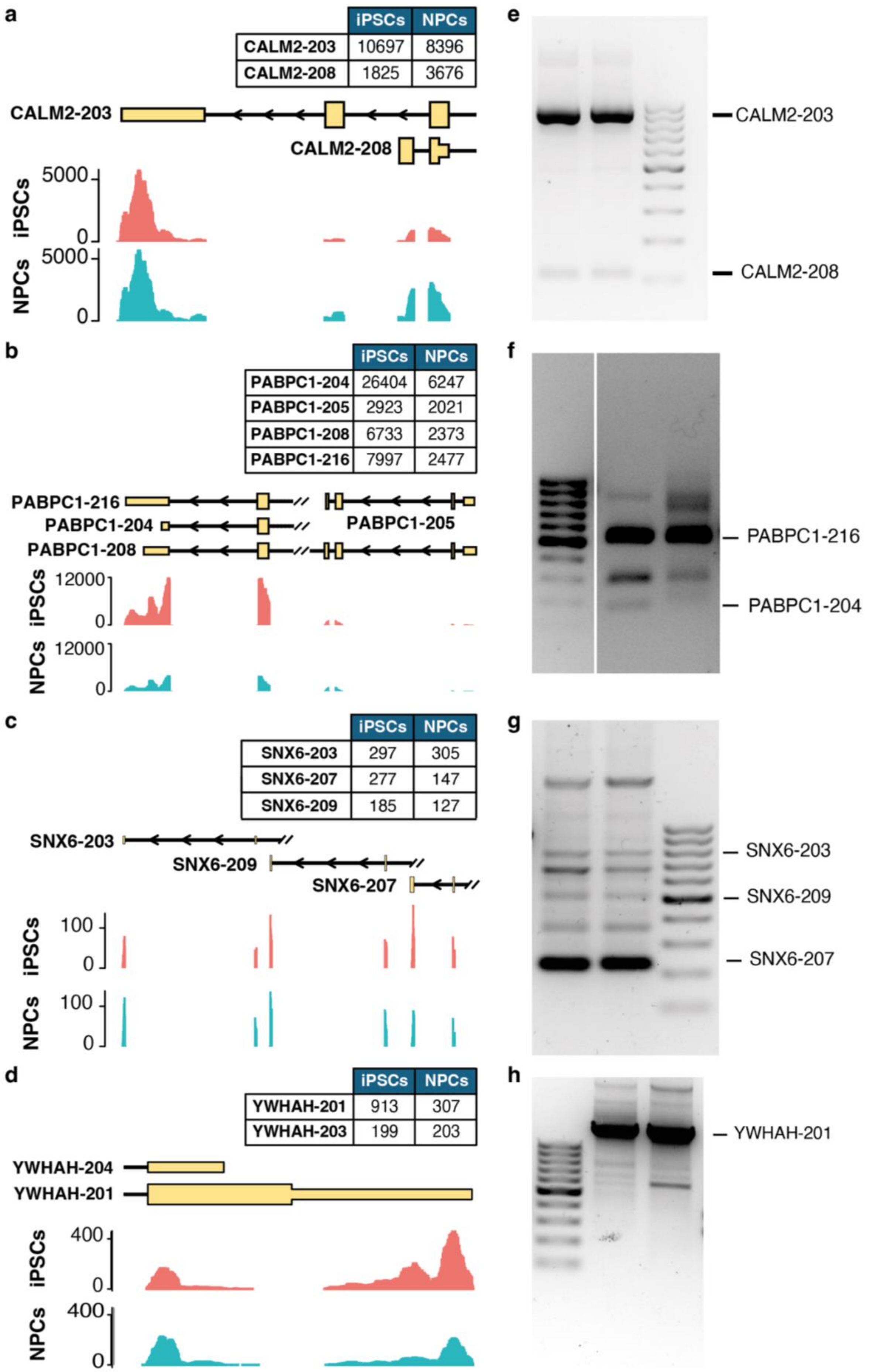

a
iPSCs NPCs
CALM2-203 10697 8396
CALM2-208 1825 3676
CALM2-203
CALM2-208
5000
iPSCs
0
5000
NPCs
0
b
iPSCs NPCs
PABPC1-204 26404 6247
PABPC1-205 2923 2021
PABPC1-208 6733 2373
PABPC1-216 7997 2477
PABPC1-216
PABPC1-204
PABPC1-205
PABPC1-208
12000
iPSCs
0
12000
NPCs
0
c
iPSCs NPCs
SNX6-203 297 305
SNX6-207 277 147
SNX6-209 185 127
SNX6-203
SNX6-209
SNX6-207
100
iPSCs
0
100
NPCs
0
d
iPSCs NPCs
YWHAH-201 913 307
YWHAH-203 199 203
YWHAH-204
YWHAH-201
400
iPSCs
0
400
NPCs
0
e
CALM2-203
CALM2-208
f
PABPC1-216
PABPC1-204
g
SNX6-203
SNX6-209
SNX6-207
h
YWHAH-201

**Figure S3. Experimental validation of isoform changes between iPSCs and NPCs. a-d.** SCALPEL isoform quantification of CALM2 **(a)**, PABPC1 **(b)**, SNX6 **(c)**, and YWHAH **(d)** genes. Coverage plots show the distribution of filtered reads along isoforms. **e-h.** Experimental validation of detected isoform for the genes in **a-d** using 3'RACE. Using this technique we detect all the isoforms of CALM2 **(e)** and SNX6 **(g)** genes and the main isoforms of PABPC1 **(f)** and YWHAH **(h)**.

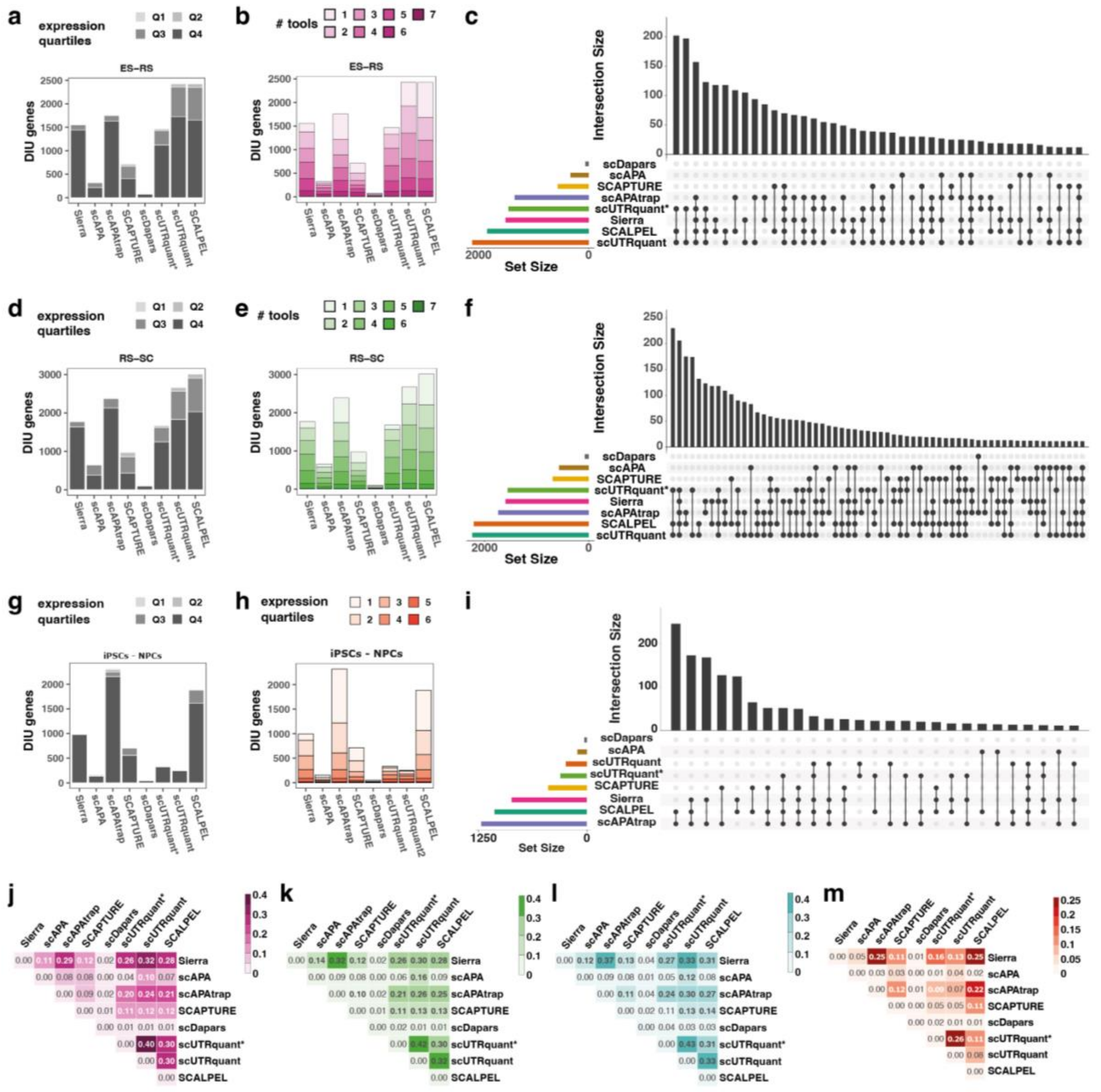


**Figure S4. Additional benchmarking of SCALPEL.** DIU genes identified by each tool between ES and RS **(a-c)**, RS and SC **(e-g)** and iPSCs and NPCs **(h-j).** SCALPEL and scUTRquant predictions are similar in the 10x datasets **(a-f)** while the number of detected DIU genes by scUTRquant drops significantly on the Drop-seq dataset **(g-i). a, d, g.** Distribution of predicted DIU genes across expression quartiles. In all comparisons, almost all genes belong to the quartiles with highly expressed genes (Q3 & Q4). **b, e, h.** Agreement in the prediction of DIU genes across tools. The agreement in the prediction of DIU genes by SCALPEL and scUTRquant using the UTRome is similar for 10x datasets **(b,e)** and superior to that of scUTRquant using the standard annotation (scUTRuant*). **c, f, I.** UpSet plots showing the agreement in the predictions from each tool between ES and RS **(c),** RS and SC **(f)** and iPSCs and NPCs **(j).** Only sets containing mmore than 10 genes are shown**. j-m.** Heatmaps comparing the agreement in the predictions across all tool pairs in ES and RS **(j),**

RS and SC **(k),** ES and SC **(l)** and iPSCs and NPCs **(m).** Heatmaps Scale bars show the Jaccard index coefficients for all comparisons. In all cases, SCALPEL and scUTRquant agreement with other tools is similar in the 10x datasets **(k-m)** and higher for SCALPEL in the Drop-seq dataset **(m)**.

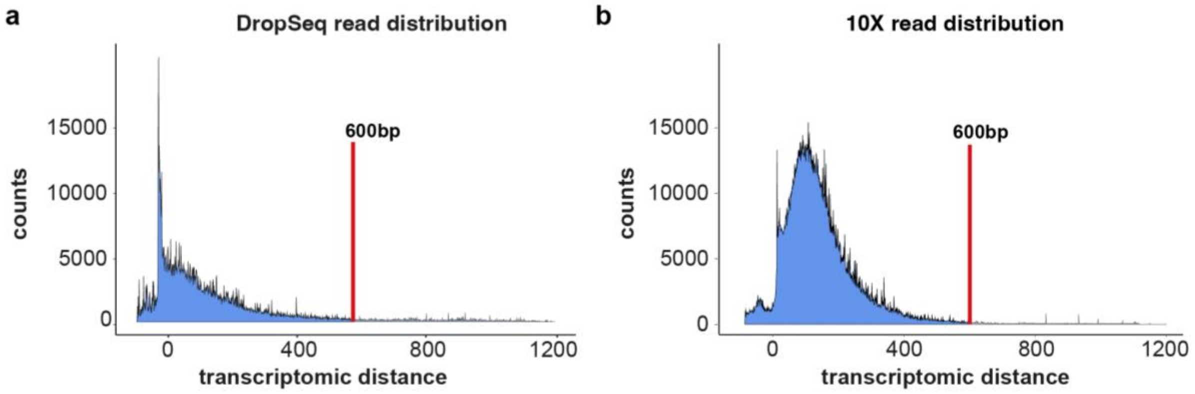


**Figure S5. Distance distribution of reads to annotated 3'ends of genes with only one expressed isoform.** Empiric distribution of the 3'end of mapped reads in the transcriptomic space relative to isoform 3' ends in the drop-seq dataset **(a)** and the 10x dataset **(b)**. These distributions are used in SCALPEL to define the probability of a read to come from a particular isoform.
